# Validation of manifold-based direct control for a brain-to-body neural bypass

**DOI:** 10.1101/2022.07.25.501351

**Authors:** E. Losanno, M. Badi, E. Roussinova, A. Bogaard, M. Delacombaz, S. Shokur, S. Micera

## Abstract

Brain-body interfaces (BBIs) are neuroprostheses that can restore the connection between brain activity and body movements. They have emerged as a radical solution for restoring voluntary hand control in people with upper-limb paralysis. The BBI module decoding motor commands to actuate the limb from brain signals should provide the user with intuitive, accurate, and stable control. Here, we present the design and demonstration in a monkey of a novel brain decoding strategy based on the direct coupling between the activity of intrinsic neural ensembles and output variables, meant to achieve ease of learning and long-term robustness. We identified once an intrinsic low-dimensional space (called manifold) capturing the co-variation patterns of the monkey’s neural activity associated to reach-to-grasp movements. We then tested the animal’s ability to directly control a computer cursor using cortical activation along the manifold axes and demonstrated rapid learning and stable high performance over 16 weeks of experiments. Finally, we showed that this brain decoding strategy can be effectively coupled to peripheral nerve stimulation to trigger hand movements. These results provide evidence that manifold-based direct control has promising characteristics for clinical applications of BBIs.

## 1. Introduction

Brain-body Interfaces (BBIs) are neuroprostheses that allow users to voluntarily control the movement of their body through an artificial neural bypass. A survey of patients with tetraplegia due to spinal cord injury [1] showed that BBIs are the preferred solution compared to the control of external robotic devices characterizing classic brain-machine interfaces (BMIs) [2]. In BBIs, brain activity recorded from motor cortical areas using invasive [3]–[10] or non-invasive [11], [12] interfaces is translated into motion commands to actuate limbs via electrical stimulation of neuromuscular structures. Thus, BBIs need to tackle two complex neurotechnological modules, i.e., a motor decoding module and a movement restoration module, and their integration [13].

Focusing on the restoration of hand function, an ideal BBI should effectively integrate an easy-to-learn, accurate, and stable brain decoding paradigm with a motor restoration module allowing the selective control of the hand. Recently, we demonstrated in a preclinical study in monkeys that peripheral nerve stimulation (PNS) at the intrafascicular level can evoke multiple grasps and hand extension movements with only two nerve implants [14], thus complying with the requirement of movement selectivity. Here, we present a brain decoding module based on the direct linear coupling between intrinsic neural ensemble dynamics and motion commands, which satisfies the characteristics of ease of learning and temporal stability. We next validate a full BBI integrating this brain decoding approach with intrafascicular PNS to trigger hand movements.

To design our brain decoding strategy, we built on recent studies [15]–[17] showing that neural population dynamics is constrained by the brain circuitry in a low-dimensional space, i.e., the neural manifold, spanned by the so-called neural modes, and that learning a new task is facilitated when the underlying neural activity pattern lies within this intrinsic manifold [17]. We hypothesized that by directly linking the activation of intrinsic neural modes to the controlled variables, the subject could learn to modulate this activation in such a manner that reduces the need for frequent calibration. Thus, we extended the previously validated approach of direct control based on the voluntary modulation of single-neuron activity aided by biofeedback [3] to the use of intrinsic neural ensemble dynamics.

We examined the performance of the manifold-based direct control strategy in a macaque monkey. Specifically, we computed once a 2D manifold capturing a significant portion of the variance of the animal’s neural activity while performing a behavioral grasping task. We then coupled the activation of the two fixed neural modes to the 2D movement of a cursor and tested this BMI paradigm in a point-to-point task with incremental variations over weeks. This BMI phase was used to evaluate the intuitiveness and long-term performance of our decoding strategy. We show that the monkey could succeed rapidly and robustly over time. Finally, we additionally coupled the dynamics of the two neural modes to the amplitude of stimuli delivered by intrafascicular electrodes implanted in the animal’s arm nerves. We demonstrate that our decoding strategy can be integrated with intrafascicular PNS into a BBI to grade hand movements.

## 2. Results

We tested a manifold-based direct control paradigm to control two degrees of freedom (DoFs) in a macaque monkey implanted with a 48-channel intracortical array in the hand region of primary motor cortex (M1). We distinguish three phases of the experimental protocol: (i) a calibration phase, in which the 2D neural manifold was identified, (ii) a BMI phase, in which the monkey used the activation of the neural modes spanning the manifold found in (i) to directly control a cursor on a screen, and (iii) a BBI phase in which the monkey used the same manifold-based direct control strategy to actuate the hand via intrafascicular PNS.

### 2.1. Calibration of a 2D brain control space based on motor neural modes

We identified an intrinsic 2D neural manifold associated with a hand motor task as the brain control space for direct control of 2 DoF cursor and hand movements. During the calibration session, we recorded M1 activity of the monkey while performing center-out reaching and grasping of objects mounted on a robotic arm [18] (**Figure 1A**). Using principal component analysis (PCA) [15], we derived the three main neural modes, representing the directions of highest variance (13%, 8%, and 5%, respectively) of the recorded M1 activity. We then examined the dynamics of the three neural modes, i.e., the so-called latent variables [15], during the motor task, to select the two control signals for the subsequent direct control experiments. We relied on the hypothesis that the two intrinsically most modulated latent variables would provide a larger working range when directly coupled to output commands. A higher modulation depth was observed for the second (mean±std across trials equal to 179±45 a.u.) and third (115±30 a.u.) latent variables with respect to the first (69±29 a.u.). Thus, we selected the 2D manifold defined by the second and third neural modes as the brain control space. The matrix mapping M1 activity into the 2D manifold was kept fixed for the rest of the experimental protocol and no other calibration session was performed.

**Figure 1.**
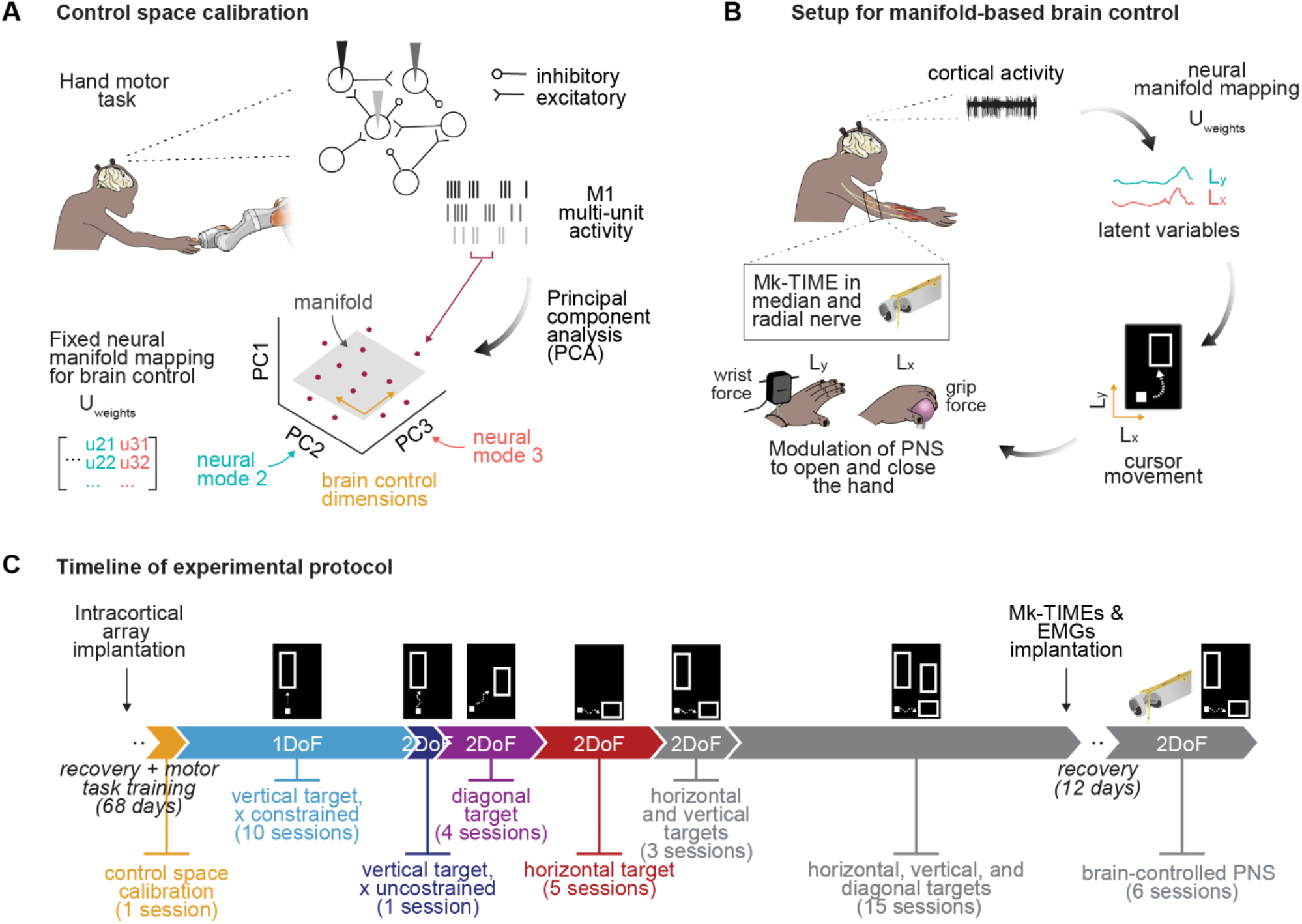
Experimental protocol for 2D manifold-based direct control. **A** Calibration of the brain control space based on neural modes, illustrated in a simplified, conceptual way with three recording channels. We applied principal component analysis (PCA) to M1 multi-unit activity recorded while the animal was performing a hand motor task and evaluated the neural space defined by the three main PCs (neural modes). The firing rate of each channel at each time instant is a point (red dot) in this space. We chose the 2D manifold (grey plane) defined by the second and third neural modes (orange arrows) as the control space for subsequent brain control experiments. The *U*_*weights*_ matrix contains the coefficients of the second and third PCs. **B** Setup for manifold-based direct control. The monkey drove a cursor (white square) in 2D (orange arrows) to reach a target box (empty rectangle) by modulating its cortical activity. The cortical activity was projected in the manifold-based control space by multiplying the firing rate of M1 channels to the *U*_*weights*_ matrix. The neural dynamics along the second and third neural modes (i.e., the second and third latent variables *L*_*y*_ and *L*_*x*_), thus computed, were linearly mapped to the cursor vertical (y) and horizontal (x) coordinates, respectively. In a second phase, *L*_*y*_ and *L*_*x*_ were also linearly linked to the stimulation amplitude of two intrafascicular electrodes implanted in the radial and median nerves, respectively, to evoke hand opening and closing. **C** Timeline of experimental protocol. The different phases of brain control experiment are depicted, i.e., the number of DoFs that the monkey had to control and the position of the target to reach with the cursor.

### 2.2. BMI with manifold-based direct control

Next, we tested the effectiveness and robustness of a 2D BMI with manifold-based direct control over 38 sessions (spanned over 113 days, **Supp. Table 1**). The monkey controlled a cursor on a screen through its M1 activity mapped into the 2D manifold (**Figure 1B**). The second and third latent variables, hereafter referred to as *L*_*y*_ and *L*_*x*_, were proportionally converted into the vertical (y) and horizontal (x) coordinates of the cursor, respectively. We designed a delayed point-to-point cursor control task: the animal had to first keep the cursor in a baseline position for 0.5 s and then reach and hold a target location for 0.1 s. Trial timeout was set to 8 s and successful trials were rewarded with liquid food. We employed an incremental training paradigm [19]: the number of DoFs to be controlled and the reaching space were progressively changed during the protocol (**Figure 1C**). For the first 10 sessions, only the y-coordinate of the cursor was brain-controlled with targets placed vertically with respect to the baseline position (cyan in **Figure 1C**): during these sessions the x-coordinate was set to 0. Next, and for the rest of the protocol, we allowed the monkey to control the cursor both in the x and y directions and we varied the location of the target: on session 11 we only presented vertical targets (blue), on sessions 12 to 15 the targets were placed diagonally to the baseline position (purple), and on sessions 16 to 20, horizontally (red). Finally, between sessions 21 and 38, the targets were randomly alternated (gray).

The monkey was able to effectively modulate its latent neural activity to perform the different tasks (**Figure 2A**). Importantly, the control was possible without using hand muscle contractions (**Supp. Figure 1**). The performance was high since day 1 of the first control configuration (1 DoF, vertical target), with 82% successful trials (**Figure 2B**) which were executed in a median time of 2.41 s (**Figure 2C**), and 21% first attempt successes (defined as the trials in which the cursor was held at the baseline and target positions for the required timespans on the first time these positions were reached) (**Supp. Figure 2A**). Over the next sessions with this configuration, we observed, despite some dips, an overall increase in success rate up to 90% on session 10 (**Figure 2B**), a significant decrease in execution time (p<0.001, F-test; **Figure 2C**), and a significant increase in the percentage of trials completed on the first attempt (p<0.01, F-test; **Supp. Figure 2A**), indicating that with practice the animal learned to perform the task more efficiently. After the introduction of the horizontal DoF, the accomplishment of the vertical target task was slightly compromised and the success rate decreased to 84% (session 11; **Figure 2B**). Another difficulty was encountered when the monkey had to jointly modulate the two latent variables. Indeed, we observed a further drop in the success rate on session 12 when the diagonal target was introduced (80% successes; **Figure 2B**). However, with training, the percentage of successful trials gradually increased, reaching 90% (session 15; **Figure 2B**) as it was reached at the end of the 1 DoF vertical task phase. Meanwhile, the execution time declined significantly (p<0.01, F-test) until a median value of 1.27 s (session 15; **Figure 2C**). When on session 16 we introduced the horizontal target, the success rate decreased to 84% (**Figure 2B**) and both the execution time (median of 2.33 s; **Figure 2C**) and the percentage of trials completed on the first attempt (25%, **Supp. Figure 2A**) returned to values close to those on the first days of the protocol. Nevertheless, over time, we observed an improvement in all these performance measures (**Figure 2B-C, Supp. Figure 2A**). After gradually adapting to the different tasks, the monkey was able to effectively switch between them. Indeed, when on session 21 we started to alternate different targets, she succeeded in 94% of the trials (**Figure 2B**) in a median time of 1.45 s (**Figure 2C**). The performance remained quite stable until session 38 (90% successes, **Figure 2B**; median execution time of 1.61 s, **Figure 2C**), corresponding to 113 days after the calibration of the control space (**Supp. Table 1**). This performance plateau possibly reflects the saturation of both the animal’s neuromodulation ability and motivation.

**Figure 2.**
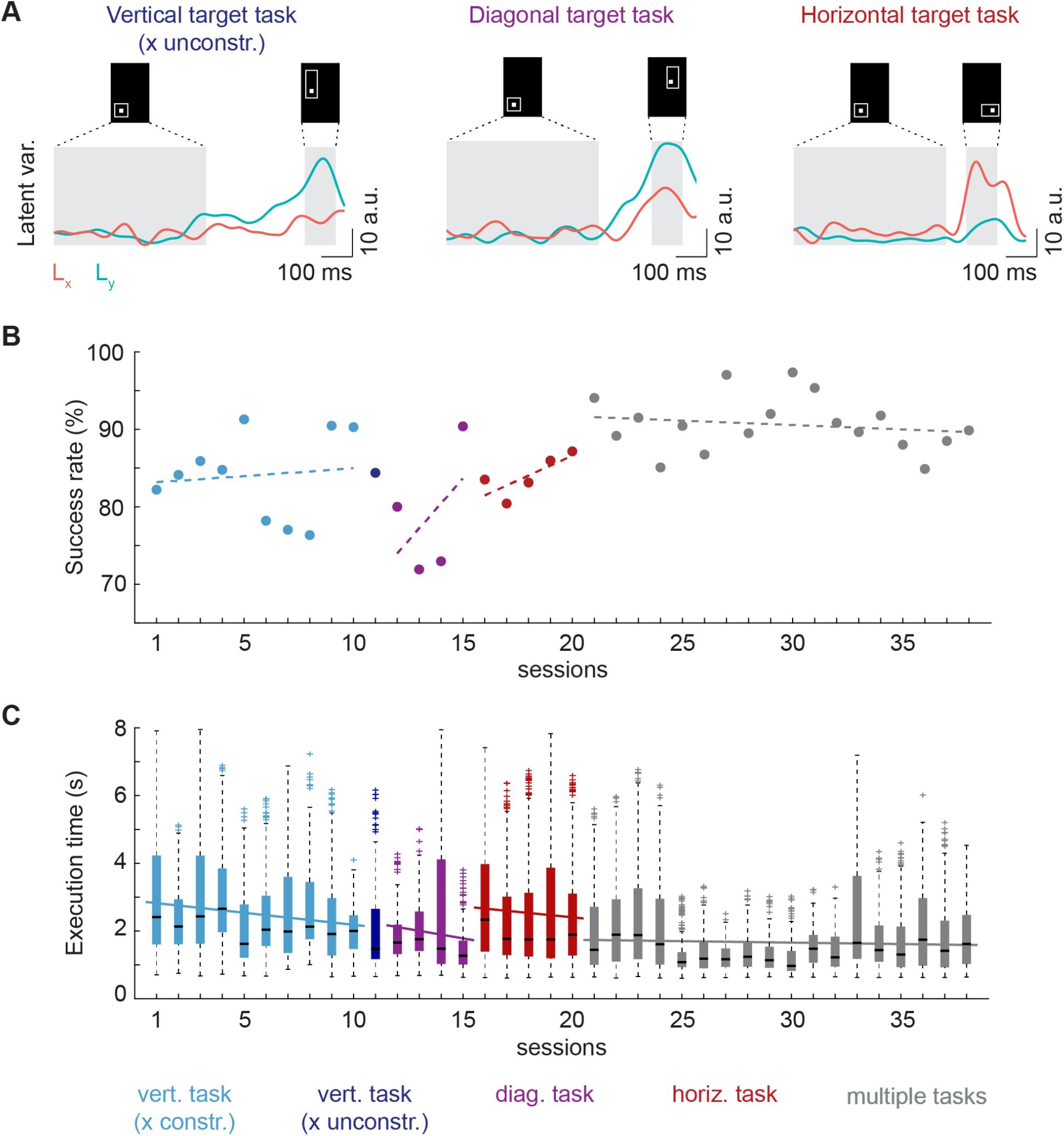
Performance of manifold-based BMI. **A** Activation of the latent variables *L*_*x*_ and *L*_*y*_ (linearly mapped to the cursor x and y coordinates, respectively) during representative successful trials of 2D cursor control for the three types of task (vertical, horizontal, and diagonal target). The task consisted in (i) maintaining the cursor in a baseline box for 0.5 s, (ii) steering the cursor toward the target box and holding it inside it for 0.1 s. The task had to be completed within 8 s for the monkey to succeed. **B** Success rate over sessions. **C** Execution time of successful trials over sessions, after outliers removal. In panels **B** and **C** the different colors indicate the different types of task performed by the animal throughout the protocol. Linear regression models were fitted to the data over the sessions with the same task (full line when significant, i.e., p<0.05, F-test, dashed line otherwise).

For the 2 DoF control configurations, we measured the movement error, i.e., the average deviation of the cursor path from the ideal straight trajectory between the baseline and target positions. Because we did not impose the path to reach the target, the monkey often succeeded in the task by exploiting curved trajectories due to the activation of both *L*_*x*_ and *L*_*y*_ for all the target types. The movement error decreased slightly over time for the horizontal target and stagnated over the multiple-target sessions (**Supp. Figure 2B**), the monkey having reached a stable success rate and execution time.

### 2.3. Neural tuning strategies

During the extended timespan of the cursor control experiment, we observed day-to-day neural recording instabilities, in agreement with previous studies [20]–[22]. Indeed, the average firing rate of M1 channels in the baseline condition (**Supp. Figure 3A**) and the corresponding latent neural activity (**Supp. Figure 3B**) varied across sessions. We thus investigated whether, following these instabilities, the animal changed its neural tuning strategy to perform the different tasks. In particular, we analyzed the inter-session variability of M1 channels preferential tuning, as measured by normalized modulation depth (see Materials and Methods), for the three targets within and between two phases of the experimental protocol, i.e., when the target of interest was the only one presented, and when it was alternated with the other targets. As expected, we observed some levels of variability in channel-wise modulation across sessions within the same protocol phase (median of 0.76, 1.07, 0.62 a.u. in the single-task phase and of 0.80, 0.81, 0.74 a.u. in the multi-task phase for the vertical, diagonal, and horizontal targets, respectively; **Figure 3A**). Interestingly, the variation between sessions of the single and multi-task phases was higher than the variation across sessions within the same phase for all the three targets (median of 1.32, 1.30, 0.78 a.u.; **Figure 3A**) and to a greater extent for the vertical and diagonal targets. This suggests that the circumstances of the task contributed significantly to the changes in neural tuning. We next analyzed the average neural tuning strategy at the single channel level in each protocol phase (**Figure 3B**) and focused on the most modulated channels (**Supp. Figure 3C**). We can see that during 1D control with only vertical targets, the monkey preferentially modulated channels #3, 13, 16, 20, and 22, all of which had a positive weight on *L*_*y*_. When the horizontal DoF was introduced, channel #20 was abandoned, likely because of its similar positive contribution to both neural modes. Moreover, the animal started to tune channel #29, associated with a positive weight on *L*_*y*_ and a slightly negative weight on *L*_*x*_, and, interestingly, channel #27, associated with a much higher weight on *L*_*x*_ than on *L*_*y*_, likely to counteract the strong negative effect of channel #22 on *L*_*x*_ and thus keep the horizontal displacement at zero. The diagonal target in the single task phase was attained by favorably tuning channels #3 and 13, which had a more positive impact on *L*_*y*_ than on *L*_*x*_, and channel #20. When introduced, the horizontal target was reached by mostly modulating channels #25 and 27, which had a much higher weight on *L*_*x*_ than on *L*_*y*_, and channel #20. These three channels were maintained in the multi-task phase of the protocol for the horizontal and diagonal targets, with slightly different ratios between each other, accompanied by channel #10, associated with a similar low positive weight on both neural modes. Channels #10, 20 and 27 were also among the most modulated to reach the vertical target in the multi-task phase, together with channels #22 and 29, which had a positive weight on *L*_*y*_ and a negative weight on *L*_*x*_ and thus were used to neutralize the movement of the cursor along x. This strategy probably proved to be the most efficient for the animal to switch between tasks. All together these results indicate that the monkey adapted its neural tuning over time led by a combination of changes in neural recordings and experimental conditions.

**Figure 3.**
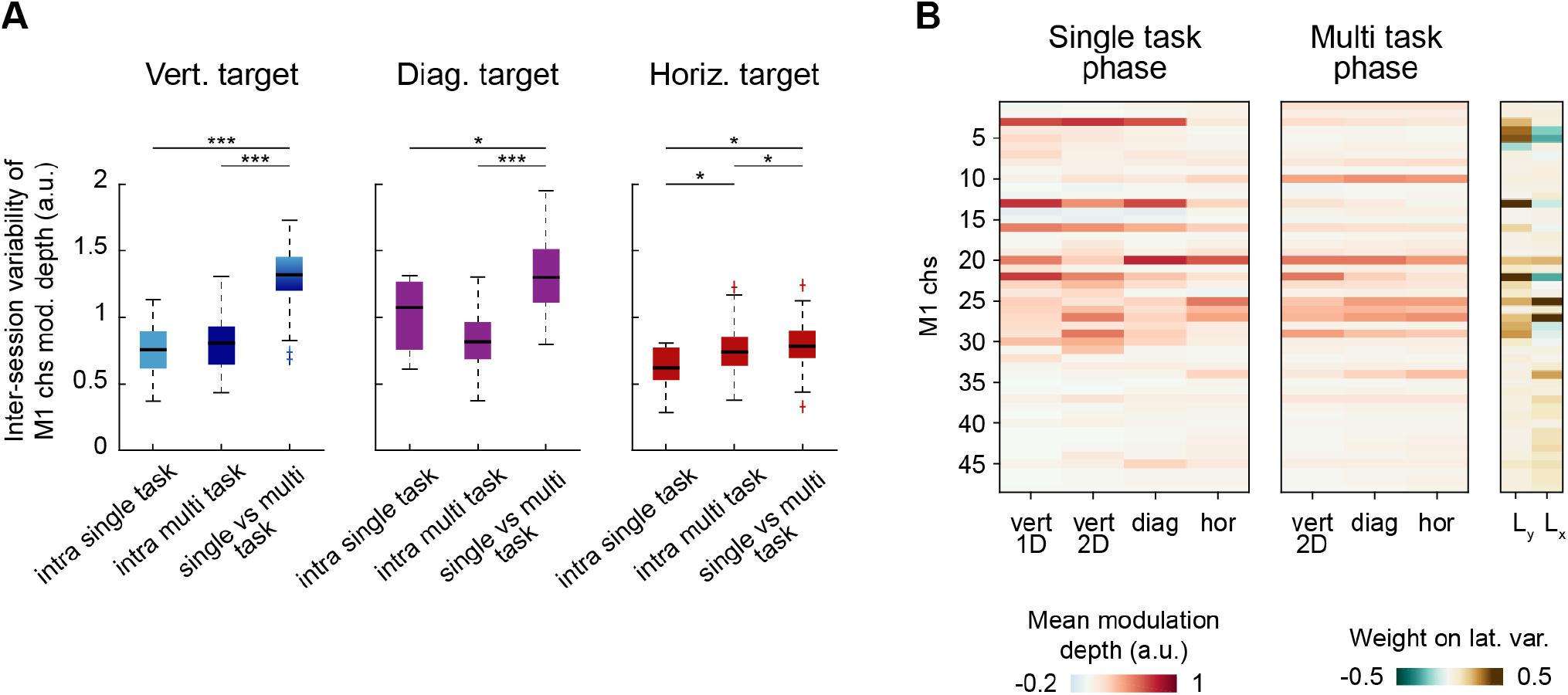
Neural tuning strategies. **A** Inter-session variability of M1 channels modulation depth within and between the two phases of the experimental protocol (i.e., single task and multi task phases) for each target type. For the vertical target, only the first 10 sessions with 1D control were considered in the single task phase, while session 11 with 2D control was excluded. **B** Normalized modulation depth of M1 channels, averaged over all trials of each protocol phase with the same target. The contribution weights of M1 channels on the two latent variables L_x_ and L_y_ are shown on the right. * p < 0.05, ** p < 0.01, *** p < 0.001, Wilcoxon rank-sum test.

### 2.4. BBI with manifold-based direct control

Finally, we investigated the feasibility of using our direct manifold-based brain control paradigm to drive a neuroprosthesis based on intrafascicular PNS and grade hand movements. For this experiment, the animal was implanted with two customized intrafascicular electrodes (Mk-TIMEs) [14], one in the median nerve and one in the radial nerve, to trigger the opening and closing of the hand, respectively. We designed the BBI experiment as follows. While the monkey performed the cursor control task with vertical and/or horizontal targets, the latent variables *L*_*x*_ and *L*_*y*_, once over a threshold, linearly modulated the amplitude of the stimuli applied to the median and radial nerve, respectively (**Figure 1B**), either jointly or independently (**Supp. Table 2**). Through a short calibration phase at the beginning of the experimental session, we set the saturation level and threshold for stimulation of the driving latent variable/s (**Supp. Figure 4A**). This latter value was regulated to reduce target-unspecific stimuli due to the frequent coactivation of *L*_*x*_ and *L*_*y*_, and at the same time span a large range of neuromodulation. The calibration also served to determine the functional amplitude range for the selected Mk-TIME channels (**Supp. Figure 4B**). After setting the control parameters, we tested the BBI in grading the two target motor functions, i.e., hand opening and closing. The full BBI protocol is described in **Figure 4A**. M1 activity was processed in real-time to extract spike events and compute the channels firing rate. Stimulation-induced artifacts were then removed by subtracting the firing rate of a channel that responded only when stimuli were applied. Noise-free spike rates were projected into the 2D manifold to derive the activation of the two latent variables. After being smoothed, *L*_*x*_ and *L*_*y*_ were linearly transformed into cursor coordinates and, in addition, the leading latent variables of the session, if over the threshold, were converted into amplitude of stimulation. Charge-balanced pulses with the defined intensity were finally applied to the nerve at a frequency of 50 Hz. The overall decoding procedure induced a time delay of approximately 10 ms. We repeated this experiment over 6 sessions.

**Figure 4.**
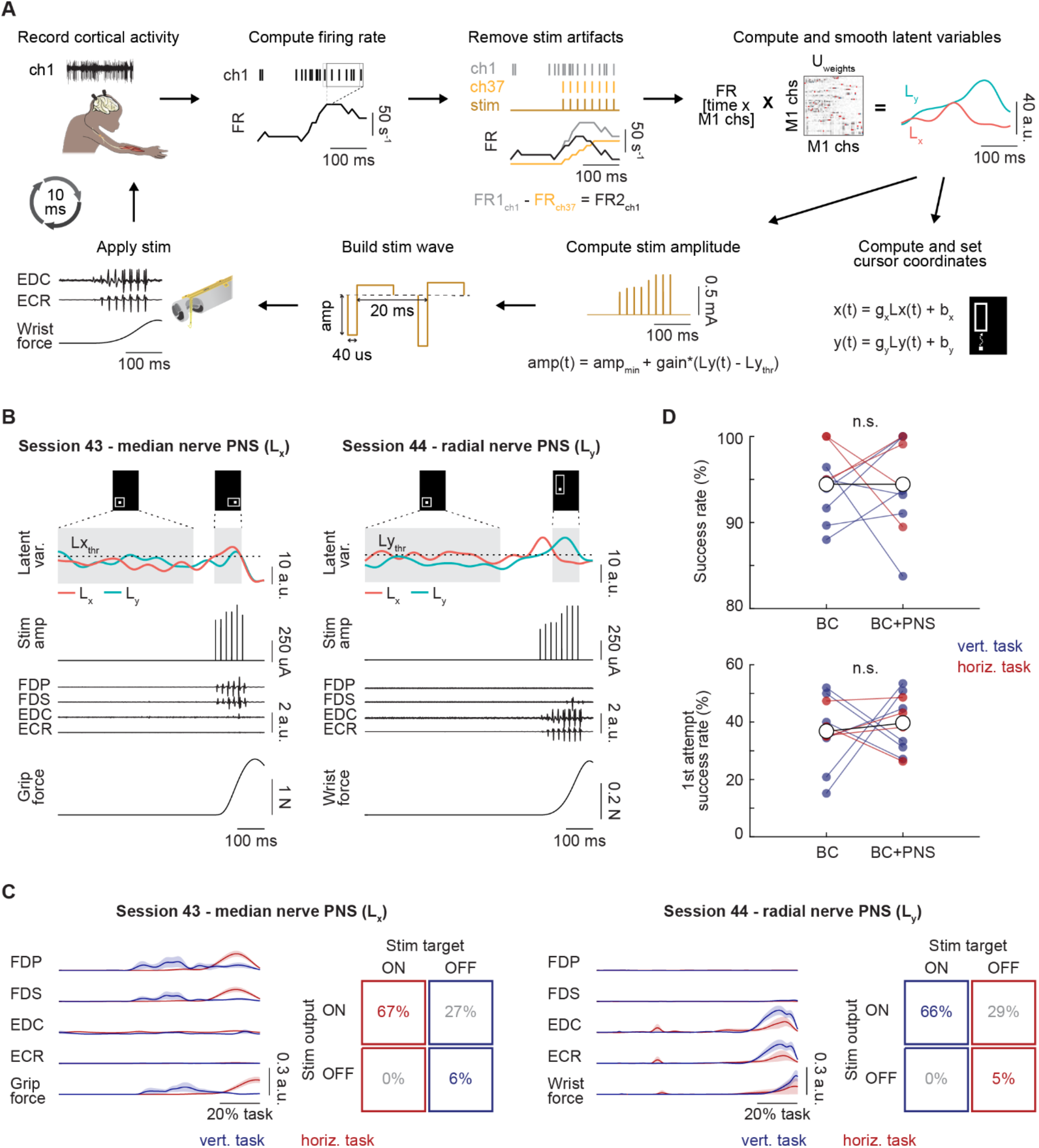
Methods and performance of manifold-based BBI. **A** Procedure for brain PNS control. M1 cortical activity is recorded. The firing rate of each M1 channel is computed as the number of spikes in overlapping bins of 100 ms with a sliding window of 10 ms. Stimulation artifacts are removed by subtracting the firing rate of a reference M1 channel (ch37), found to respond only when stimulation was applied. The latent variables *L*_*x*_ and *L*_*y*_ are computed by multiplying the firing rate of the 48 M1 channels per the *U*_*weights*_ matrix. After being smoothed, *L*_*x*_ and *L*_*y*_ are linearly transformed to set the cursor x and y coordinates. The leading latent variables of the session are also linearly mapped to the amplitude of PNS (in the example, only *L*_*y*_ is driving stimulation). The stimulation wave is built as a train of biphasic pulses (pulse-width of 40 us, frequency of 50 Hz). Stimulation is then applied from the preselected channel (in the example of the radial Mk-TIME) thus recruiting hand muscles and generating force. The overall decoding procedure induces a time delay of approximately 10 ms. **B** Representative successful trials of brain PNS control on two sessions in which median or radial nerve stimulation was enabled, respectively. The monkey performed the brain cursor control task while the latent variables controlled both the movement of the cursor and the amplitude of PNS. On session 43, *L*_*x*_ was linearly mapped to median nerve stimulation to recruit flexor muscles and close the hand. On session 44, *L*_*y*_ was linearly mapped to radial nerve stimulation to recruit extensor muscles and open the hand. Stimulation was enabled only after succeeding in the baseline phase of the cursor control task and activated when the leading latent variable exceeded the threshold. **C** Quantification of the target specificity of the BBI on session 43 (only median nerve stimulation enabled, controlled by *L*_*x*_) and on session 44 (only radial nerve stimulation enabled, controlled by *L*_*y*_). Left for each session: hand muscle activity and force, generated by stimulation, averaged across all successful trials with the same target type (vertical and horizontal). Right for each session: confusion matrices showing the percentage of successful trials in which stimulation was activated or kept off as desired or not (i.e., median nerve stimulation, which was controlled by *L*_*x*_, should ideally have been delivered for the horizontal target and kept off for the vertical target, whereas radial nerve stimulation, which was controlled by *L*_*y*_, should ideally have been delivered for the vertical target and kept off for the horizontal target). **D** Comparison of success rate in the brain cursor control task when PNS was or was not enabled (n = 10, 6 sessions). Data referring to the same session and the same target type (vertical or horizontal) were pairwise compared. On the 6 sessions, median nerve stimulation was modulated by *L*_*x*_, whereas radial nerve stimulation was modulated by *L*_*y*_. Except on the first session, only one type of stimulation was enabled at a time (**Supp. Table 2**). n.s. = not significant (p>0.05, Wilcoxon signed-rank test). Abbreviations: flexor digitorum profundus (FDP), flexor digitorum superficialis (FDS), extensor digitorum communis (EDC), extensor carpi radialis (ECR).

*L*_*x*_-driven median nerve stimulation effectively activated the hand flexors to smoothly close the hand and modulate grip force when the animal accomplished the horizontal target task (**Figure 4B** left). Conversely, *L*_*y*_-driven radial nerve stimulation recruited the hand extensors to incrementally open the hand and grade wrist extension force during the vertical target successes (**Figure 4B** right). In the two sessions in which the same type of stimulation (i.e., median, or radial) was enabled for both targets (**Supp. Table 2**), we quantified the target specificity of the BBI. In 27% of the successful trials on session 43, *L*_*x*_ exceeded the stimulation threshold during the vertical target task, inducing spurious median nerve stimuli and hand flexor responses (**Figure 4C** left). Similarly, in 29% of the successes on session 44, *L*_*y*_ exceeded the threshold during the horizontal target task, undesirably triggering the radial nerve and the hand extensors (**Figure 4C** right). Thus stimulation was not selectively activated in the majority of the cases, even though the undesired motor responses had a minor strength compared to those desired (**Figure 4C**).

We next controlled if PNS perturbed the brain cursor control task. Considering all six sessions, we did not observe a significant decrease in either the success rate (p=0.38, Wilcoxon signed-rank test) nor the percentage of trials completed on the first attempt (p=0.25, Wilcoxon signed-rank test) compared with the PNS-free setting (**Figure 4D**), confirming the efficacy of our procedure for stimulation artifacts removal.

## 3. Discussion

We assessed the performance of a 2-DoF brain control strategy confined within a fixed intrinsic motor manifold [15] for novel BBI restoring hand movements. We employed a simple yet intuitive brain decoding module based on a direct linear coupling between latent neural dynamics and output commands.

First, we assessed the within-manifold neuromodulation ability of the monkey in a 2D delayed point-to-point cursor control task. This BMI paradigm provided us with the flexibility necessary to study the long term temporal and task-related effects on the decoder performance. Our brain control strategy proved to be easy-to-learn and robust over 16 weeks. The animal showed a high success rate from the first day of the experiment without prior training and adapted readily to new tasks. We did observe small drops in proficiency when a change in neuromodulation strategy was required, but these were easily compensated for with little practice. Compared to previous studies in which monkeys were exposed to 2D cursor control based on a fixed linear decoder applied to a stable ensemble of neurons [23], [24], learning was more rapid. This result was certainly favored by the incremental design of the training protocol [19], but is also likely due to the “ecological” BMI mapping employed. By fixing the control space within an intrinsic manifold, we exploited natural (i.e., already acquired) neural activity patterns [17], and by intuitively relating the cursor movement to these patterns, we facilitated learnability. The monkey was then able to consistently switch between the different tasks, maintaining a success rate of ∼90% until the end of the protocol (113 days after the control space calibration, **Supp. Table 1**). This long-term robustness is a promising result, as neural recording instabilities in chronic settings constitute one of the main challenges for the clinical translation of BMIs [20]–[22]. Standard BMIs based on algorithms to decode movement-related parameters from neuronal population activity require ad-hoc unsupervised decoder-updating methods [25]–[28] to account for day-to-day changes in neural recordings and avoid the frequent collection of calibration data. On the other hand, decoders that rely on stable single neurons [3] or stable neuronal ensembles [23] have limited temporal applicability because the isolation of the same cells is disrupted by neural turnover, which happens after a period of days to weeks [23]. Here, as expected, we did observe changes in neural recordings over the study period. However, the monkey was able to adjust the tuning of neural ensembles, also depending on the experimental conditions, to consolidate its skills in the different tasks and preserve a high success rate over several weeks. We believe that this effortless adaptation is still due to the inherence of manifold-based control. These results expand previous findings on the potential and utility of neural plasticity for BMI applications [23], [24].

As a final step, we conducted a pilot experiment to test our direct manifold-based control strategy in driving a PNS-based neuroprosthesis for grading hand opening and closing. By training the monkey to timely up-regulate latent neural activity that linearly modulated the amplitude of intrafascicular PNS, our approach enabled the timely triggering of smoothed hand movements. Importantly, although it certainly elicited sensory percepts [29], the stimulation of healthy nerves did not impair performance in cursor control. These proof-of-concept results demonstrate the feasibility of integrating our decoding paradigm into a BBI.

A limitation of our approach was the limited accuracy in effector control. The monkey frequently reached the visual target along curved cursor paths due to activation of both latent variables. This led, in the BBI phase, to the target-unspecific application of stimuli to the median and radial nerves, resulting in weaker but frequent undesired muscle responses. Our training paradigm, which was based on a simple point-to-point cursor control task and was lacking of instructions that encouraged straight cursor trajectories, certainly did not favor accuracy. In view of applying this control strategy to motor functions that require the coordinated recruitment of hand flexors and extensors, a more constrained task, such as an instructed-path [30] or a pursuit-tracking [31] task, should be used in the future to promote independent and finer control of the latent variables. This scenario would also be crucial to investigate whether our proposed decoder can achieve the level of control accuracy and smoothness provided by state-of-the-art algorithms such as the Kalmar filter [32], [33]. Moreover, while we have limited our BBI paradigm to the control of two motor DoFs, necessitating only two driving latent variables, extending it to more complex movements will require additional control signals. In this framework, it will become increasingly critical to ensure the decoupling of latent neural dynamics to separately control multiple stimulation channels targeting specific muscles or muscle synergies. We thus note that a crucial aspect that should be investigated to corroborate the clinical utility of this approach would be to determine the degree of dominance and independence that can be achieved on multiple neural modes. Finally, further validation with a larger number of monkeys is necessary to generalize our results.

In the perspective of clinical translation to people with severe motor disabilities, some practical points need to be discussed. First, the efficacy of a manifold identification method based on imagined or attempted movements has yet to be validated. However, since M1 was shown to be amply engaged not only in overt movements but also in cognitive motor processes [34], we believe that goal-directed motor imagery or motor attempt would be effective calibration paradigms, as usual in BMI and BBI clinical applications [35]. We also point out that brain areas such as premotor or parietal cortices could provide an interesting alternative or complement to M1 to derive intrinsic low-dimensional spaces associated with motor control [15], [36]. Second, the choice of the calibration tasks may be critical for the ease-of-learning of the BBI. Here, the neural manifold was identified based on a center-out reaching movement which was structurally related to the point-to-point cursor motion. Although experimental verification of this point is lacking, our recommendation would be to select calibration tasks that are congruent with the final BBI task. In the same line, we note that a larger repertoire of calibration movements may be necessary to provide the user with greater versatility for more complex control. Third, while this approach is more directly applicable to patients suffering from motor disorders that do not affect the cerebral cortex, such as spinal cord injury or brainstem stroke, neural tuning adaptability after cortical injuries remains to be tested. Since it was shown that cortical stroke survivors can learn to modulate ipsilesional cortical rhythms [37], we believe that control of latent neural dynamics is also possible, and could be enhanced by brain stimulation [37]. Moreover, studies have shown that BBIs can promote neurological recovery [37]–[42] thanks to the contingent link between brain activity and body mobilization which triggers Hebbian-like plasticity [43]. Therefore, we believe that our BBI would act like a reinforcing loop that simultaneously exploits and promotes neural plasticity.

We conclude that direct control based on latent neural dynamics is a promising paradigm for BBI control in clinical applications because of its reliability and long-term stability, resulting from the inherence of neural manifolds and the intuitiveness of direct control links.

## 4. Materials and Methods

### 4.1. Animal and implants

The experiments were conducted on an adult female *Macaca fascicularis* monkey (5 years old, 3.1 kg). The experimental protocol was elaborated in compliance with the national law on animal protection and approved by the Federal and local veterinary authorities (authorization number 2017_03_FR).

During a first surgical intervention, the monkey received the implantation of three 48-channel microelectrode arrays (Blackrock Microsystems, USA, 400 µm pitch, 1.5 mm tip length). One array was implanted in the hand region of the M1 of the right hemisphere. Primary somatosensory and premotor cortices were also implanted but not analyzed in this study. Almost 6 months later (**Supp. Table 1**), the animal underwent a second surgery. Two custom-made chronic intrafascicular multichannel electrodes (TIMEs) tailored to the monkey anatomy (Mk-TIMEs) [14] were inserted into the animal’s median and radial nerves, which innervate most of the flexor and extensor muscles of the hand, respectively [14]. The median Mk-TIME was implanted ∼2 cm proximally to the elbow and the radial Mk-TIME ∼2 cm proximally to the epicondyle along the humeral bone. In addition, to record EMG activity, the monkey was chronically implanted with 8 pairs of Teflon-coated stainless steel wires in the following flexor and extensor muscles of the hand: flexor carpi radialis (FCR), palmaris longus (PL), flexor digitorum profundus (FDP), flexor digitorum superficialis (FDS), extensor carpi radialis (ECR), extensor digitorum communis (EDC), extensor carpi ulnaris (ECU) and abductor pollicis longus (APL). The two surgeries were performed under aseptic conditions and general anesthesia induced with midazolam (0.1 mg/kg), methadone (0.2 mg/kg), and ketamine (10 mg/kg) and maintained under continuous intravenous infusion of propofol (5 ml/kg/h) and fentanyl (0.2-1.7 ml/kg/h).

### 4.2. Experimental setup and procedure

#### 4.2.1. Behavioral reach-and-grasp task

The monkey was trained to perform a center-out reach-and-grasp task, which is detailed in [18]. Briefly, a robotic arm (Intelligent Industrial Work Assistant, IIWA – KUKA, Augsburg, Germany) with seven degrees of freedom, presented custom-molded, silicone objects of various shapes (cylindrical, spherical and small triangular) in front of the animal at different locations in space. The monkey was trained to freely reach for the object with its left hand, grasp it, and then pull it towards its body by counteracting the force exerted by the robotic arm, which increased proportionally to the horizontal displacement. A trial was considered successful if the robot end-effector passed a predetermined distance threshold. Upon success, the monkey automatically received a liquid food reward through a sipper tube.

#### 4.2.2. Identification of the 2D motor manifold

M1 cortical activity recorded over an entire session of the behavioral reach-and-grasp task (including 625 trials and all periods between trials), was used to compute the axes spanning the 2D manifold, i.e., the neural modes coefficient matrix *U*_*weights*_ (**Figure 1A**). The firing rate of each M1 channel was computed offline as the number of spikes in non-overlapping bins of 10 ms. PCA was then applied to the firing rates of the 48 M1 channels to derive the *U*_*weights*_ matrix, as follows:

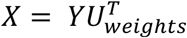

Where, *X* [*time x M*1 *chs*] is the matrix of firing rates of the 48 M1 channels, *U*_*weights*_ [*M*1 *chs x M*1 *chs*] is the matrix of PC coefficients, and *Y* [*time x M*1 *chs*] contains the PC scores, i.e., the representation of *X* in the PC space.

The *U*_*weights*_ matrix thus computed was used in the brain control experiments as a linear transformation between the firing rates and the neural activity along the main neural modes [15]:

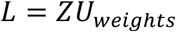

Where *Z* [*time x M*1 *chs*] is the matrix of neural firing rates of the 48 M1 channels and *L* [*time x M*1 *chs*] is the matrix of the 48 latent variables, i.e., the cortical activity projected along the neural modes. Based on their modulation depth during the behavioral motor task (see Results), we selected the second and third latent variables, hereafter referred to as *L*_*y*_ and *L*_*x*_ respectively, as control signals in the brain control experiments.

#### 4.2.3. Brain cursor control experiment

The monkey was seated in a custom primate chair in front of a large computer screen. The left arm and hand were immobilized with padded plastic restraints. Latent neural activity directly controlled a moving cursor on the screen that provided visual feedback to the animal in real-time (**Figure 1B**). Specifically, after being downsampled at 25 Hz, the two latent variables *L*_*x*_ and *L*_*y*_ were linearly transformed into the cursor horizontal (x) and vertical (y) coordinates, respectively, as follows:

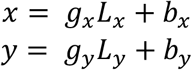

The gains *g*_*x*_ and *g*_*y*_ were manually set to 0.4 and 0.25, respectively, based on the size of the screen and the amplitude of the modulation of *L*_*x*_ and *L*_*y*_, and kept constant for the overall experimental protocol. The offset values *b*_*x*_ and *b*_*y*_ were adjusted during each session depending on the baseline neural activity, which changed across sessions likely because of changes in neural recordings (**Supp. Figure 3A-B**). The cursor was prevented from exiting the screen through boundaries on its x and y coordinates.

The task consisted in delayed 2D point-to-point cursor control. The animal had to maintain latent neural activity at a baseline level and then to up-regulate it. More precisely, at the beginning of a trial, an empty square, representing the baseline box, appeared in the lower left corner of the screen. The animal had to hold the cursor within this square for 0.5 s. Upon success in this first phase, the baseline square disappeared and a new empty rectangle appeared on the screen, in a position that differed depending on the phase of the experimental protocol (see **Figure 1C** and the “Brain cursor control timeline” section). The monkey had to move the cursor to this target box and hold it within it for 0.1 s. To succeed and thus receive a liquid food reward, the animal had to complete the overall task within 8 s. The distance between the target and the baseline boxes was set manually and varied during each session, trying to get the animal to modulate its neural activity as much as possible, but at the same time avoiding demotivating the animal.

#### 4.2.4. Brain cursor control timeline

We analyzed 38 sessions (spanned over 113 days, **Supp. Table 1**) of brain cursor control experiment, during which the monkey was gradually trained to control up to 2 DoFs and to reach different target positions (**Figure 1C**). During the first 10 sessions, the monkey had to modulate the cortical activity along only one neural mode (1 DoF control). The cursor was moved only along the y axis proportionally to the activation of *L*_*y*_ (the x coordinate was set to 0), to reach a vertical target. We then introduced the horizontal component to the cursor trajectory that was proportional to the activation of *L*_*x*_ and maintained this 2 DoF control configuration for all the subsequent sessions. For one day we presented only vertical targets, forcing the monkey to up-regulate the activity of *L*_*y*_ while maintaining the activity of *L*_*x*_ at a baseline level to succeed in the task. We then shifted the target along the horizontal axis in a diagonal position to promote the simultaneous modulation of *L*_*y*_ and *L*_*x*_. After 4 sessions, we started to present only horizontal targets to encourage the monkey to exclusively up-modulate *L*_*x*_ while keeping *L*_*y*_ at a baseline level. Once the animal achieved a success rate comparable to the other tasks (after 5 sessions), we started to randomly alternate vertical and horizontal targets and repeated for 3 sessions. The next 15 days of recordings consisted in randomly alternating vertical, horizontal, and diagonal targets.

Few sessions were excluded from the analysis because the triggers designating the task events were not properly recorded, and few others because the monkey was not motivated to perform the task as she was not in perfect health.

#### 4.2.5. Brain PNS control experiment

During the brain PNS control experiment, the animal performed the cursor control task while latent neural activity drove both the cursor movement and the stimulation amplitude of preselected channels of the median and radial Mk-TIMEs (**Figure 1B**). We selected a channel of the median Mk-TIME that recruited flexor muscles to trigger hand closure and a channel of the radial Mk-TIME that recruited extensor muscles to produce hand opening. Stimulation delivered by the median channel was controlled by *L*_*x*_, whereas stimulation applied from the radial channel was controlled by *L*_*y*_. Specifically, we modulated the amplitude *amp*(*t*) of the pulses injected through the channel of interest over time based on a linear mapping with the latent variable activation *L*(*t*), smoothed by a 100 ms moving average filter:

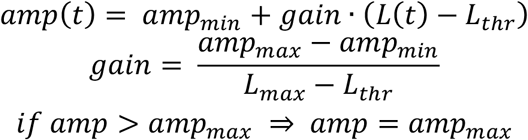

The pulse-width was fixed at 40 µs, and the frequency of the pulses at 50 Hz. The parameters of the linear relationship between *amp*(*t*) and *L*(*t*) were tuned during a calibration phase at the beginning of each session, as specified in the “Calibration of the parameters for brain PNS control” section. Stimulation was enabled only after the animal succeeded in the baseline phase of the cursor control task and disabled in between trials.

The experiment was performed for 6 sessions, in which we enabled one or both types of stimulation (i.e., median, or radial) and presented one or both types of target (i.e., vertical, or horizontal), as specified in **Supp. Table 2**.

#### 4.2.6. Calibration of the parameters for brain PNS control

At the beginning of each session of the brain PNS control experiment, a calibration procedure was performed to tune the parameters of the linear relationship between latent variable activation and stimulation amplitude (**Supp. Figure 4**). In the first step, the animal performed the brain cursor control task for approximately 10 minutes, alternating between vertical and horizontal targets (**Supp. Figure 4A**). This phase served to determine the range of latent variable modulation that the animal exhibited for the two target types on that day. Based on these recordings, we determined *L*_*max*_ and *L*_*thr*_. *L*_*max*_ of a given latent variable was set to be just above the maximum of its activation averaged across the successful trials with the target type for which it was leading (horizontal target for *L*_*x*_ and vertical target for *L*_*y*_). Conversely, *L*_*thr*_ was set to be just above the maximum of the latent variable activation averaged across the successful trials with the other target type. In this way, we aimed to exploit a wide range of neural modulation in PNS control, while limiting spurious stimuli due to a non-straight path of the cursor to the target (ideally, median nerve stimulation, controlled by *L*_*x*_, would have been activated only for horizontal targets and radial nerve stimulation, controlled by *L*_*y*_, only for vertical targets). In a second step, we applied stimulation bursts from the selected Mk-TIME channels with increasing amplitude values (pulse-width of 40 us, frequency of 50 Hz) (**Supp. Figure 4B**). In this way, we derived the amplitude range [*amp*_*min*_, *amp*_*max*_] that we used for brain PNS control. *amp*_*min*_ corresponded to the minimum amplitude at which a movement twitch occurred and *amp*_*max*_ corresponded to the amplitude at which a strong contraction movement was observed. At the end of the calibration phase, the experimenter set the calibration parameters using a graphical user interface on the control computer.

#### 4.2.7. Data acquisition

Neural signals were acquired at 30 kHz with a Neural Signal Processor (Blackrock Microsystems, USA) using the Cereplex-E headstage. Multiunit activity was thresholded (6.25x root mean square value calculated over a window of 5 s) to extract spike events. During the brain control experiments, a custom C++ routine (Visual Studio®, USA), running on a control computer, processed the neural signals in real-time to compute the latent variables. Specifically, the firing rate of each M1 channel was computed as the number of spikes in overlapping bins of 100 ms with a sliding window of 10 ms. Stimulation artifacts (only for the brain PNS control experiment) and then movement artifacts were suppressed as described in the section below. The latent variables *L*_*y*_ and *L*_*x*_ were then calculated by multiplying the noise-free firing rates of the 48 M1 channels per the *U*_*weights*_ matrix. *L*_*y*_ and *L*_*x*_ were streamed via UDP to a computer running a custom MATLAB (MathWorks, Natick MA) routine. This routine converted the latent variables into cursor coordinates, placed the visual targets on the screen, and controlled a peristaltic pump that delivered a liquid food reward. The timing of various events in the task, such as start and end of a trial, success, etc., were sent as digital triggers to the Neural Signal Processor through a synchronization board (National Instruments, US). During the brain PNS control experiment, the conversion of the latent variables into amplitude of stimulation was implemented by the C++ routine running on the control computer.

In the sessions following the implantation of the EMG electrodes, bipolar EMG signals were acquired at 12 kHz by the RZ2 processor (RZ2, Tucker David Technologies, USA) after amplification (1000×, PZ5, Tucker David Technologies, USA) using a 16-channels active headstage (LP32CH - 16, Tucker Davis Technologies, USA).

In the last two sessions of brain PNS control experiment, we measured the grip force using a custom-made sensor [18] or the wrist extension force using a commercial dual-range force sensor (Vernier, EducaTEC AG,CH) when median or radial nerve stimulation was enabled, respectively. These signals were recorded at 1 kHz using the RZ2 processor.

#### 4.2.8. Artifacts removal from neural recordings

Stimulation artifacts were removed from neural recordings by subtracting the firing rate of a reference M1 channel, found to be silent when stimulation was not applied, from the firing rate of all M1 channels.

Movement artifacts were suppressed by ensuring that if more than 40 channels (over the 128 channels of the three implanted brain arrays) had a firing rate greater than 20 spikes/s, those channels were discarded (i.e., their firing rate was set to 0).

#### 4.2.9. Electrical stimulation

Electrical stimulation was delivered through a 32-channels headstage (LP32CH - 32, Tucker Davis Technologies) using the IZ2H stimulator (Tucker David Technologies, USA) as bursts of asymmetric charge-balanced cathodic-first biphasic pulses. Stimulation waveforms were digitally built within the processor unit (RZ2, Tucker Davis Technologies) using the user programming interface OpenEx suite (Tucker Davis Technologies). Custom code was used to communicate with the controller through C++ (Visual Studio) API.

#### 4.2.10. Hand muscle activity monitoring

To show that the animal performed the brain cursor control task without exploiting hand movements, we recorded the corresponding muscle activity in two sessions after the implantation of the EMG electrodes. In these two sessions the monkey also performed the behavioral reach-and-grasp task. We compared the EMG activity of the implanted muscles acquired during the brain cursor control task with the activity measured during the behavioral task (**Supp. Figure 1**).

### 4.3. Data analysis

#### 4.3.1. Analysis of latent variables modulation during the behavioral task

To select two among the three main latent variables to be used as control signals in the brain control experiments, we computed their modulation depth during the behavioral reach-and-grasp task. We applied a 50 ms moving average filter to the latent variable activation signal and then calculated the difference between its maximum and minimum values in each motor trial.

#### 4.3.2. Analysis of changes in neural recordings and tuning

To evaluate the changes in neural recordings across sessions, we computed the mean firing rate of M1 channels during the baseline phase of the cursor control task (i.e., when the cursor was in the baseline box) and averaged across all the trials of each session (**Supp. Figure 3A**). Similarly, for each trial we computed the mean activity of latent variables *L*_*x*_ and *L*_*y*_ during the baseline phase (**Supp. Figure 3B**).

To evaluate the neural tuning strategy used by the monkey to reach the different targets, we measured the modulation depth of the 48 M1 channels. Modulation depth was computed as the difference between the channel’s maximum firing rate during the target holding phase of the cursor control task (i.e., when the cursor was in the target box) and its mean firing rate during the baseline phase. To focus on which channels were preferentially modulated rather than to what extent, the modulation depth of all channels was normalized to the maximum across channels for each trial. The neural tuning strategy of each session was considered as the 48-element vector obtained by averaging over all trials. The variability in neural tuning strategy across sessions within and between the two main phases of the experimental protocol (i.e., single-task and multi-task phases) (**Figure 3A**), was calculated as the Euclidean norm of the difference in neural tuning strategy between each pair of sessions within the same phase or between phases. The neural tuning strategy of each protocol phase (**Figure 3B**) was considered as the average over all trials of all sessions belonging to that phase. The most modulated channels (**Supp. Figure 3C**) were considered as those showing a modulation depth higher than *q*3 + *w* × (*q*3 − *q*1), where *w* is a multiplier constant, and *q*1 and *q*3 are the 25^th^ and 75^th^ percentiles of all channels data related to that phase and target. *w* was set to 1.5 for the vertical and diagonal targets, 2.5 for the horizontal target.

#### 4.3.3. Performance assessment in the brain cursor control experiment

Performance in the brain cursor control experiment was assessed by counting the percentage of successful trials and measuring the execution time and movement error. Trials were considered successful if, in less than 8 s, the monkey was able to i) hold the cursor in the baseline box for 0.5 s and ii) reach the target box and hold the cursor inside it for 0.1 s. Among the successful trials, we distinguished the successes on the first attempt, i.e., the trials in which the monkey succeeded in holding the cursor in the baseline and target boxes for the required timespans on the first time the cursor entered the respective box. The execution time of successful trials was calculated as the interval between the appearance of the baseline box on the screen and the completion of the task. The movement error was computed for successful trials as 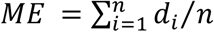[44], where *d*_*i*_ is the distance of the *i*_*th*_ point of the cursor path from the line connecting the centers of the baseline and target boxes (*d*_*i*_ ≥ 0). *ME* measures the offset of the cursor path from the ideal straight trajectory. Execution time and *ME* outliers (elements lying outside 1.5 times the interquartile range) were removed for each session. Linear regression models were fitted to the data of the described measures over the sessions the animal performed the same type of task.

#### 4.3.4. Performance assessment in the brain PNS control experiment

We evaluated the monkey’s ability to successfully perform the brain cursor control task even when PNS was enabled, by comparing the success rate obtained during the brain PNS control task with that obtained during the calibration phase on the same session. Results obtained for the same target type (vertical and horizontal) were pairwise compared.

For the two sessions in which only one PNS type was enabled for both vertical and horizontal targets (sessions 43 and 44, **Supp. Table 2**), we assessed the percentage of successful trials in which stimulation was target-selectively applied. Specifically, median nerve stimulation (session 43), which was controlled by *L*_*x*_, should ideally have been delivered for the horizontal target and kept off for the vertical target. Conversely, radial nerve stimulation (session 44), which was controlled by *L*_*y*_, should ideally have been delivered for the vertical target and kept off for the horizontal target. This analysis quantifies the monkey’s ability to modulate the two latent variables independently and also reveals the appropriateness of the chosen PNS control parameters.

#### 4.3.5. EMG and kinetic signals processing

EMG signals were band-pass filtered between 50 and 500 Hz. A Savitzky-Golay filter with a smoothing window of 2.5 ms was applied to remove stimulation artifacts. The envelope was computed by rectifying the EMG signal and applying a low-pass filter at 6 Hz. Signals were normalized to the maximal muscle activity obtained across the trials of interest.

Grip and wrist force signals were low-pass filtered at 10 Hz and detrended by subtracting a cubic spline fitted on the data outside the stimulation periods. Voltage values were converted to Newtons using the calibration curves of the two sensors [18], (Vernier, EducaTEC AG,CH). Signals were normalized to the maximum across the trials of interest.

#### 4.3.6. Statistics

Data are reported as mean *±* standard error of the mean (s.e.m.) unless specified otherwise. Statistical significance of linear regression models was evaluated using the F-test. Statistical significance of the difference between two samples was evaluated using the non-parametric Wilcoxon rank-sum test for unpaired data and the non-parametric Wilcoxon signed-rank test for paired data.

## Acknowledgements

The authors would like to thank Prof. Eric M. Rouiller and Dr. Marco Capogrosso for helpful advice and discussions; Dr. Sophie Wurth and Dr. Simon Borgognon for help with the experiments; J. Maillard and L. Bossy for the care provided to the monkey; A. Zbinden for the veterinary survey of the animal. Funding for this study was provided by the Swiss National Science Foundation grant NeuGrasp (205321_170032), the Wyss Center for Bio and Neuroengineering, and the Bertarelli Foundation.

## Supplementary Figures

**Supplementary figure 1.**
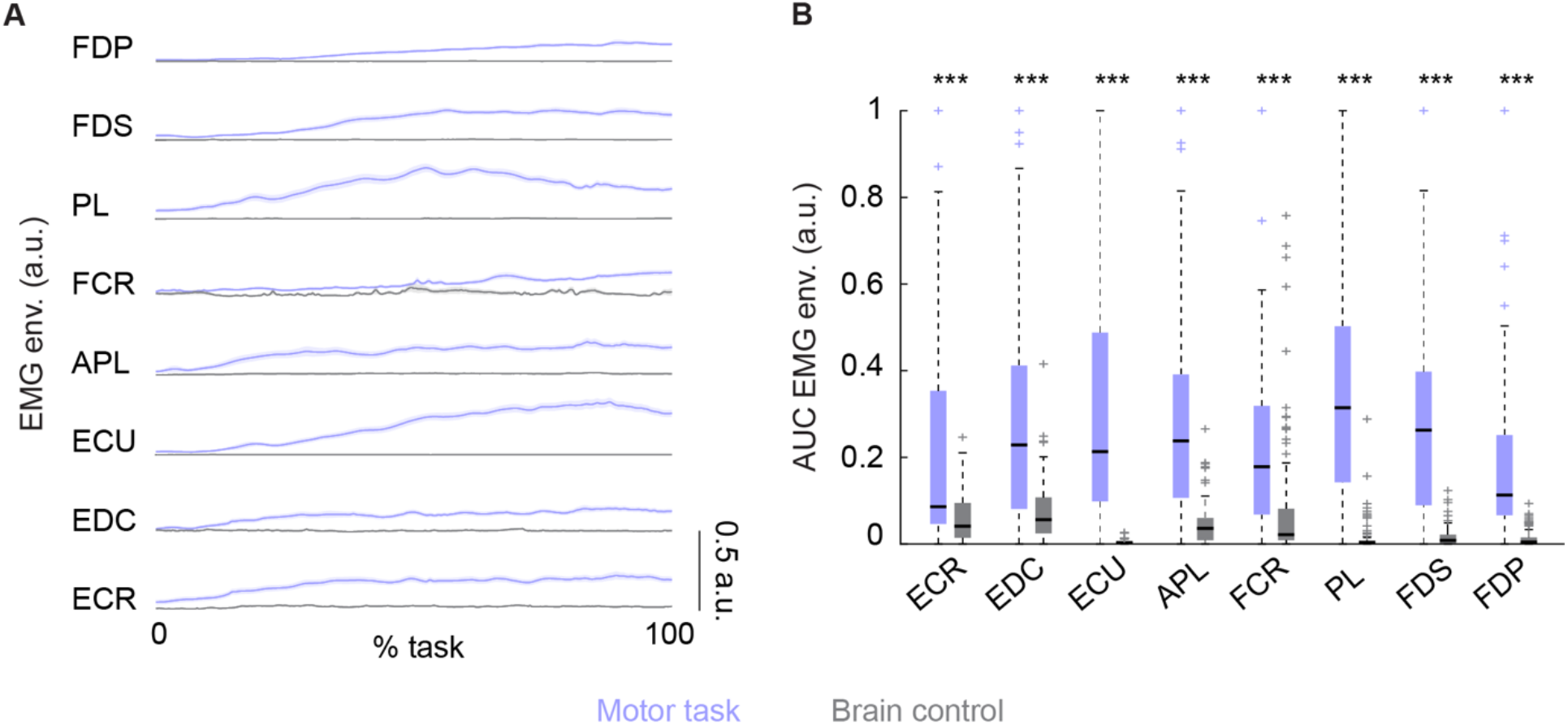
Hand muscles activity during manifold-based BMI and hand motor tasks. **A** Average dynamics of the EMG envelope of hand muscles in comparison between reach-and-grasp trials and brain cursor control trials (session 39). **B** Area under the curve (AUC) of the EMG envelope for the different muscles in comparison between reach-and-grasp trials and brain cursor control trials (sessions 39 and 43). *** p < 0.001, Wilcoxon rank-sum test. Abbreviations: flexor digitorum profundus (FDP), flexor digitorum superficialis (FDS), palmaris longus (PL), flexor carpi radialis (FCR), abductor pollicis longus (APL), extensor carpi ulnaris (ECU), extensor digitorum communis (EDC), extensor carpi radialis (ECR).

**Supplementary figure 2.**
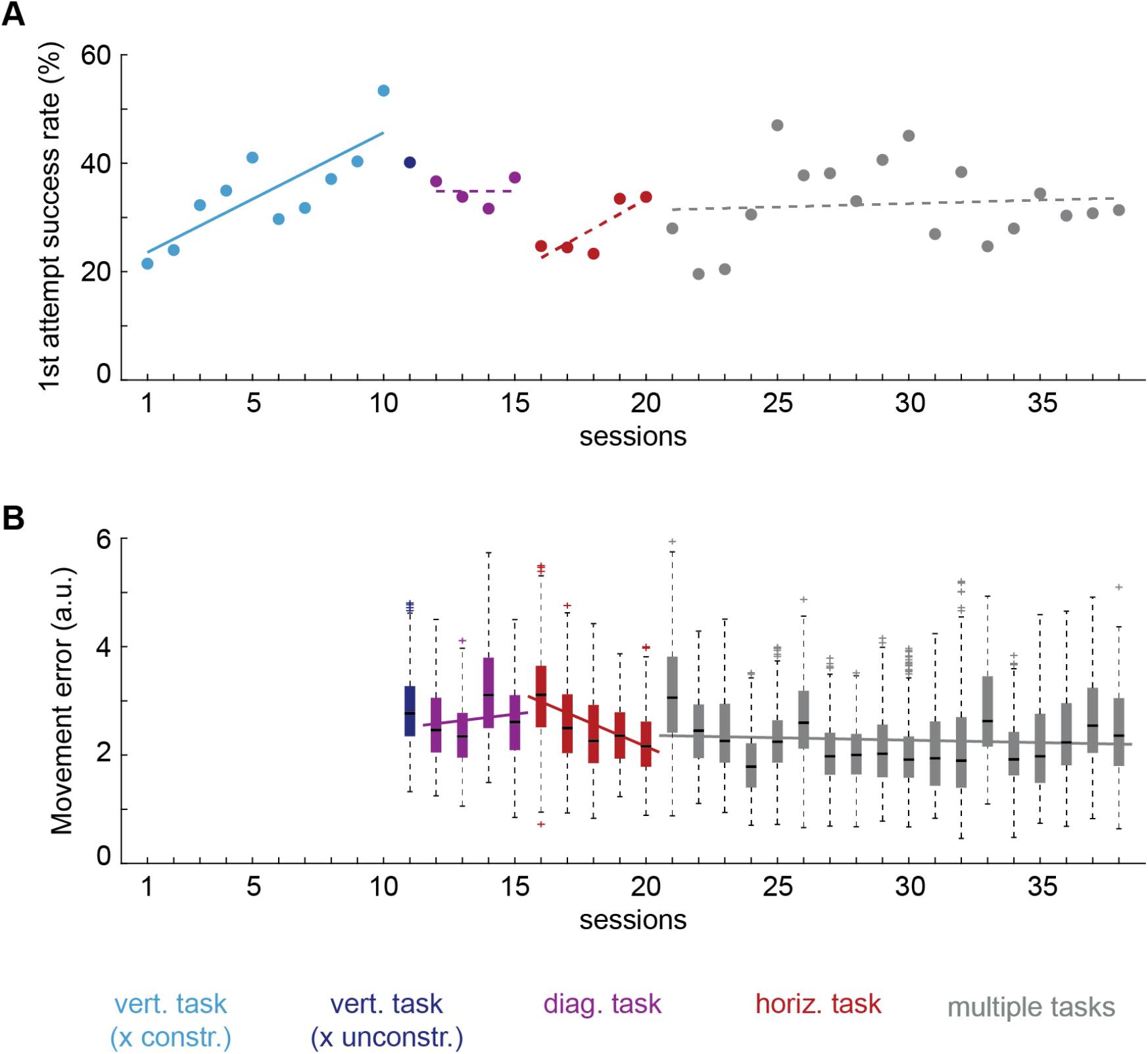
Additional performance measures of manifold-based BMI. **A** Percentage of 1^st^ attempt successes (i.e., the monkey holds the cursor in the baseline and target boxes for the required periods on the first time the cursor entered each box) over sessions. **B** Movement error (i.e., deviation of the cursor path from the ideal straight trajectory connecting the centers of the baseline and target boxes) of successful trials over sessions, after outliers removal. For the 1 DoF configuration of the first 10 sessions, the movement error could not be computed. In the two panels, the different colors indicate the different types of task performed by the animal throughout the protocol. Linear regression models were fitted to the data over the sessions with the same task (full line when significant, i.e., p<0.05, F-test, dashed line otherwise).

**Supplementary figure 3.**
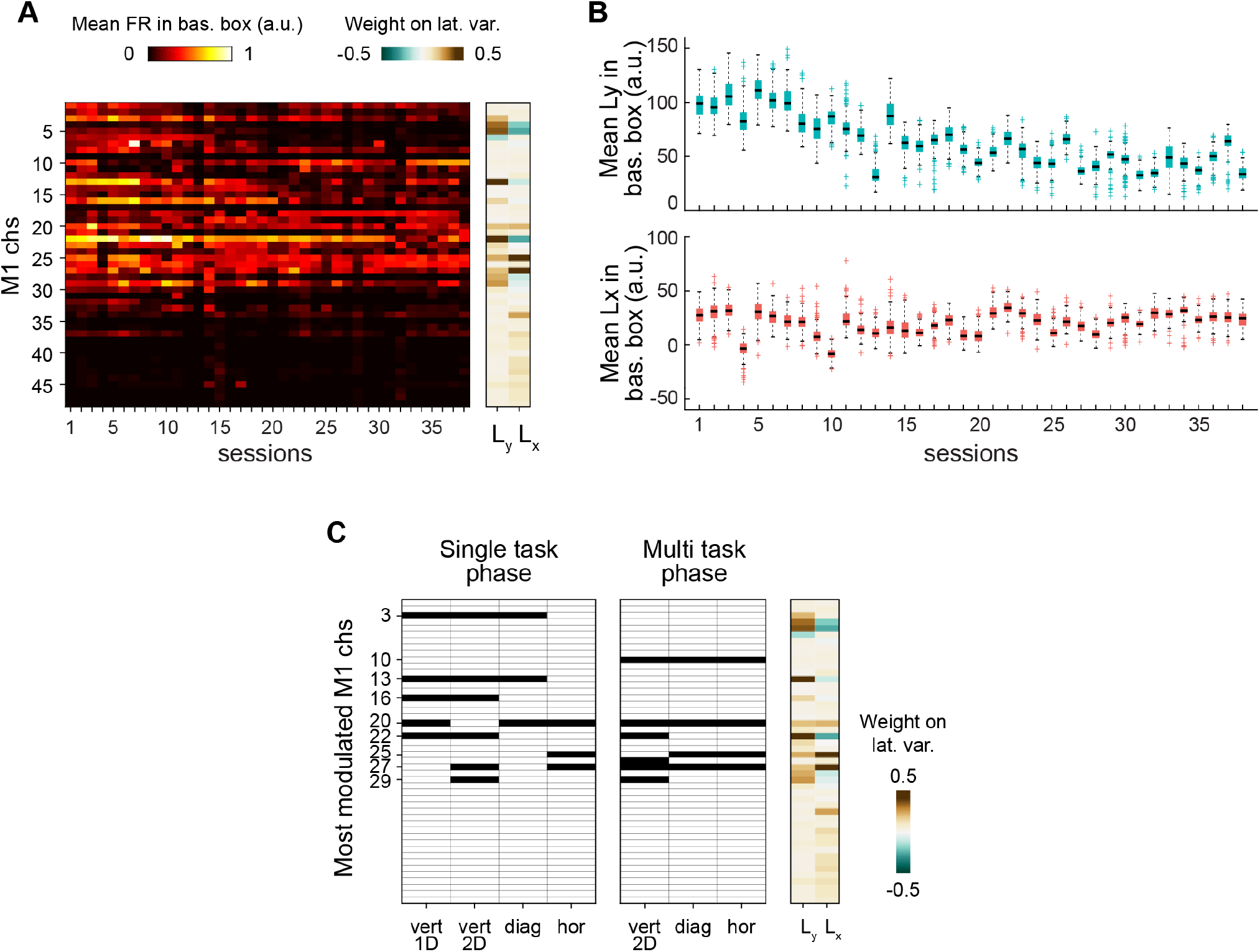
Changes in neural recordings and tuning across sessions. **A** Mean firing rate of M1 channels during the baseline phase of the cursor control task (i.e., when the cursor was in the baseline box) across sessions. The contribution weights of M1 channels on the two latent variables L_x_ and L_y_ are shown on the right. **B** Mean latent variables L_x_ and L_y_ during the baseline phase of the cursor control task across sessions. **C** Most modulated M1 channels (see Supp. Methods) for each target in the two protocol phases (i.e., single task and multi task phases), marked in black. The contribution weights of M1 channels on the two latent variables L_x_ and L_y_ are shown on the right.

**Supplementary figure 4.**
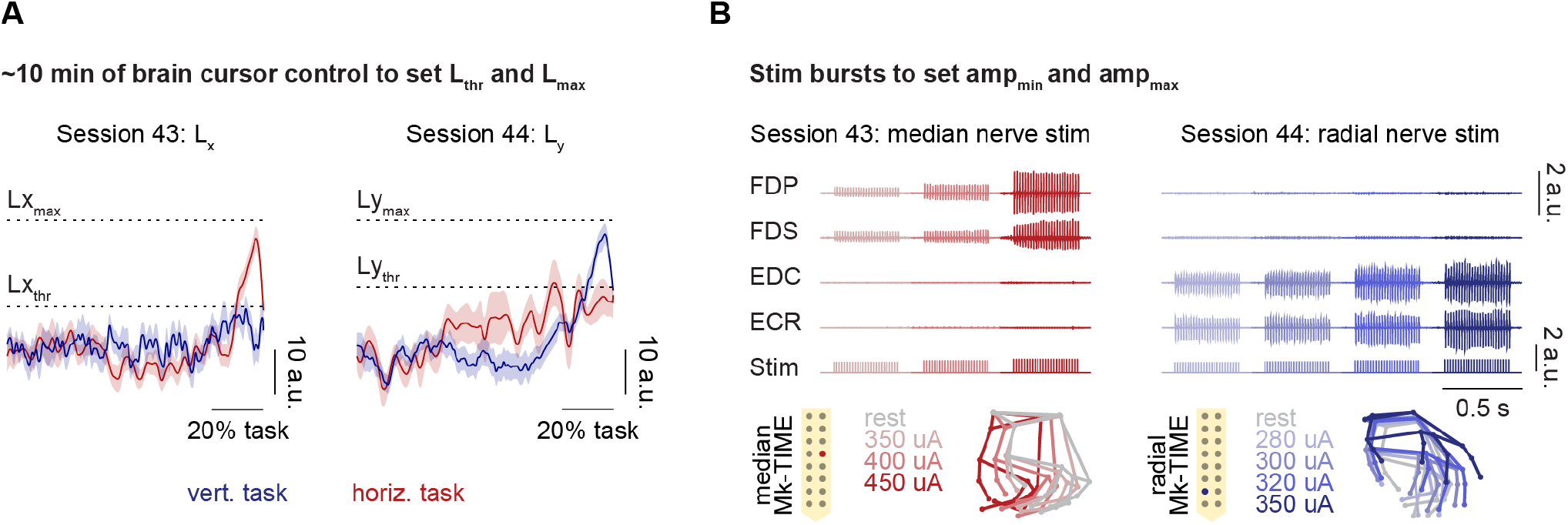
Calibration of the parameters for brain PNS control. **A** First, the animal performs the brain cursor control task for ∼10 min, alternating between vertical and horizontal targets. *L*_thr_ and *L*_*max*_ are set based on the activation of the leading latent variable/s of successful trials with the two target types (see Supplementary Methods). Two sessions are shown: session 43 (only *L*_*x*_ controlled PNS) and session 44 (only *L*_*y*_ controlled PNS). **B** Second, stimulation bursts are applied from the preselected channel of the median and/or the radial nerve with increasing amplitude values. *amp*_*min*_ and *amp*_*max*_ are set based on the motor response (see Supplementary Methods). The same two sessions as before are shown: session 43 (the monkey brain-controlled PNS applied from a channel of the median Mk-TIME that evoked hand closing) and session 44 (the monkey brain-controlled PNS applied from a channel of the radial Mk-TIME that evoked hand opening).

## Supplementary Tables

**Supplementary Table 1.**
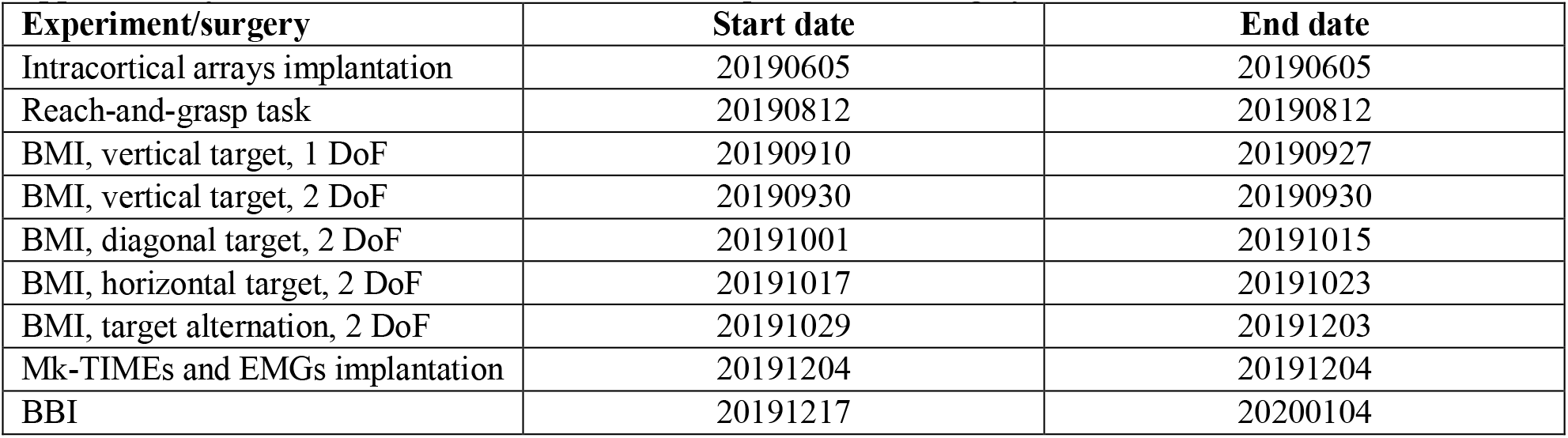
Start and end dates of each experiment and surgery.

**Supplementary Table 2.**
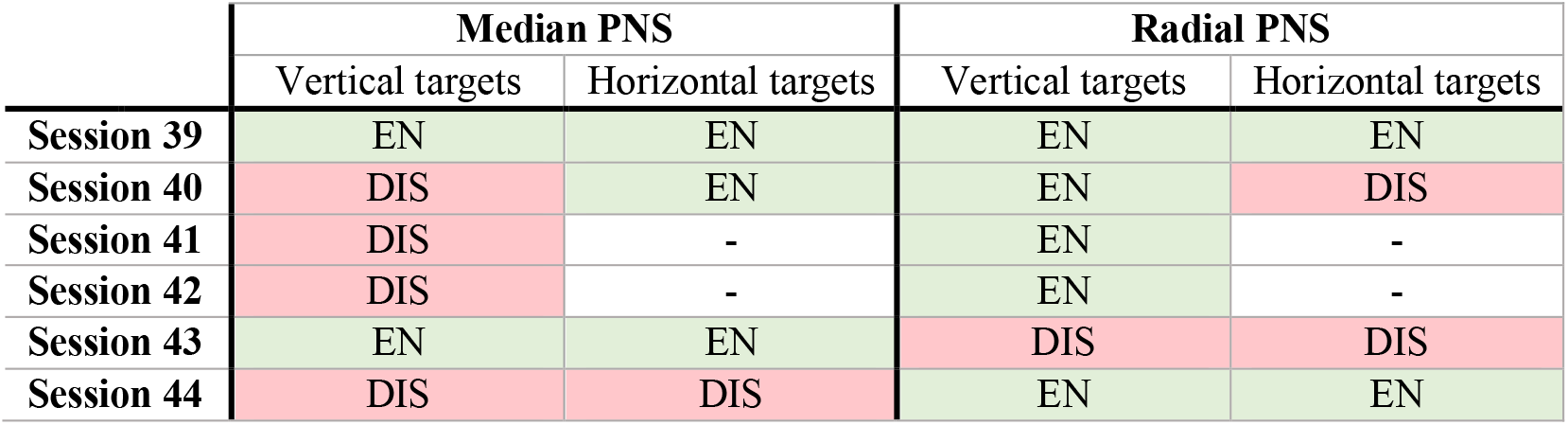
Type of stimulation enabled for the two target types on the 6 sessions of brain PNS control. Abbreviations: enabled (EN), disabled (DIS), target not presented (-).

